# More evidence for a long-latency mismatch response in urethane-anaesthetised mice

**DOI:** 10.1101/837690

**Authors:** Jamie A. O’Reilly, Thanate Angsuwatanakul

## Abstract

Prolonged electrophysiological responses to oddball stimuli have recently been observed from anaesthetised rodents. This deviant-related activity is found to extend through 200 to 700 ms post-stimulus; a window typically obstructed from analysis by the response to subsequent stimuli in the auditory sequence. A simple methodological development in terms of difference waveform computation using two adjoining evoked responses has enabled visualisation of this activity over a longer window of analysis than previously available. In the present study, the double-epoch subtraction technique was retroactively applied to data from 13 urethane-anaesthetised mice. Oddball paradigm waveforms were compared with those of a many-standards control sequence, confirming that oddball stimuli evoked long-latency potentials that did not arise from standard or control stimuli. Statistical tests were performed at every time point from 0 to 700 ms post stimuli to highlight regions of significant difference. Oddball-induced mismatch responses were found to display significantly greater long-latency potentials than identical stimuli presented in an equal-probability context. As such, it may be concluded that long-latency potentials were evoked by the oddball condition. How this feature of the anaesthetised rodent mismatch response relates to human mismatch negativity is unclear, although it may be tentatively linked to the human P3a component, which is considered to emerge downstream from mismatch negativity.

## 1. Introduction

Human mismatch negativity (MMN) is isolated from the difference waveform between auditory evoked potential (AEP) responses to stimuli presented in an oddball paradigm. This is computed by subtracting the AEP to a frequently repeated standard stimulus from that of a rarely presented deviant stimulus; i.e. *MMN* = *deviantAEP* − *standardAEP*. The MMN component is found within this difference waveform at approximately 100 to 250 ms post stimulus onset (Näätänen et al., 2012). This electrophysiological component is important clinically as a potential biomarker for neuropsychiatric disease (Baldeweg and Hirsch, 2015), spurring considerable research activity in developing rodent models of MMN (Harms et al., 2016). These may be referred to more generally as mismatch responses. Rodents are widely believed to share similar neural circuitry responsible for generating MMN in humans. Like humans, rodent mismatch responses are present while animals are conscious or anaesthetised (Nakamura et al., 2011).

There is some evidence that a long-latency mismatch response (>200 ms post stimulus) can be elicited from urethane-anaesthetised rodents (Casado-Román et al., 2019; Chen et al., 2015; O’Reilly, 2019). Curiously, there was an example of this in the first published mismatch negativity study in anaesthetised rats, elicited by the deviant-alone control paradigm (Ruusuvirta et al., 1998), which hitherto has not garnered much attention in the preclinical mismatch response literature. The majority of proposed rodent analogues of MNN have been reported to fall within the latency range of 50 to 200 ms (Nakamura et al., 2011). The deviant-alone control may actually reflect an exaggerated oddball condition (O’Reilly, 2019). Multi-unit activity recorded from prefrontal areas in urethane-anaesthetised rats appears to support this notion (Casado-Román et al., 2019). In this study, deviant stimuli triggered an increase in spiking activity from 200 to 700 ms which was absent from several control conditions; importantly, this long-latency response was also elicited by deviant-alone stimuli. This is a comparatively underinvestigated latency range of the rodent mismatch response, because it is typically obscured from analysis by the response to subsequent stimuli in the auditory sequence. However, a double-epoch subtraction may be applied to visualise deviant-evoked activity over an extended window of analysis (O’Reilly, 2019).

This paper presents a retrospective analysis of data from a mismatch negativity study in urethane-anaesthetised mice (Kurkela et al., 2018). The double-epoch subtraction method was used to examine the resulting difference waveforms across a longer period than was previously considered. This is designed to establish whether a long-latency mismatch response was elicited by oddball stimuli in this experiment.

## 2. Materials & Methods

### 2.1. Data

The data under analysis was obtained from a previous study (Kurkela et al., 2018). The original manuscript describes the conditions of data collection in detail. To summarise essential points for the current article, epidural field potentials were recorded from 13 urethane-anaesthetised mice using a wire electrode positioned above the left auditory cortex. Cerebellar referencing and a ground connection on the animal’s neck were used, and auditory stimuli were presented to the right ear. Electrophysiology signals were band-pass filtered between 1 and 30 Hz. The oddball paradigm included 4 kHz standards, with 3.5 kHz and 4.5 kHz oddballs; all were 50 ms pure sinusoids with 5 ms tapering. This may be referred to as a “balanced” oddball paradigm because it contains two oddball stimuli which deviate by equal amounts in opposite directions from the standard. This can be useful for establishing whether there inherent effects of the physical parameter being manipulated. Two versions of the oddball paradigm were presented: one with an inter-stimulus interval (ISI) of 350 ms, and another with an ISI of 600 ms. The many-standards control included physically identical stimuli to those employed in the oddball paradigm among 16 total different frequency stimuli ranging from 3.3 to 4.8 kHz in 100 Hz increments; each presented at the same rate as deviant stimuli in the oddball paradigm with an ISI of 375 ms.

### 2.2. Waveform Analysis

The double-epoch subtraction was performed on waveforms from oddball and control paradigms, as described previously (O’Reilly, 2019). In the case of oddball paradigms, *standard:standard* AEP pairs were subtracted from *deviant:standard* AEP pairs to view deviant-evoked activity over a larger window than would otherwise be available for analysis. The same operation was performed on data from the control paradigm, although it may be noted that the second AEP in each pair was produced by the average of a random selection of different frequency stimuli due to pseudo-randomness within the many-standards design. Auditory-evoked potentials and their corresponding difference waveforms are plotted from − 100 to 850 ms about the first stimulus onset time in Figure 1.

**Figure 1:**
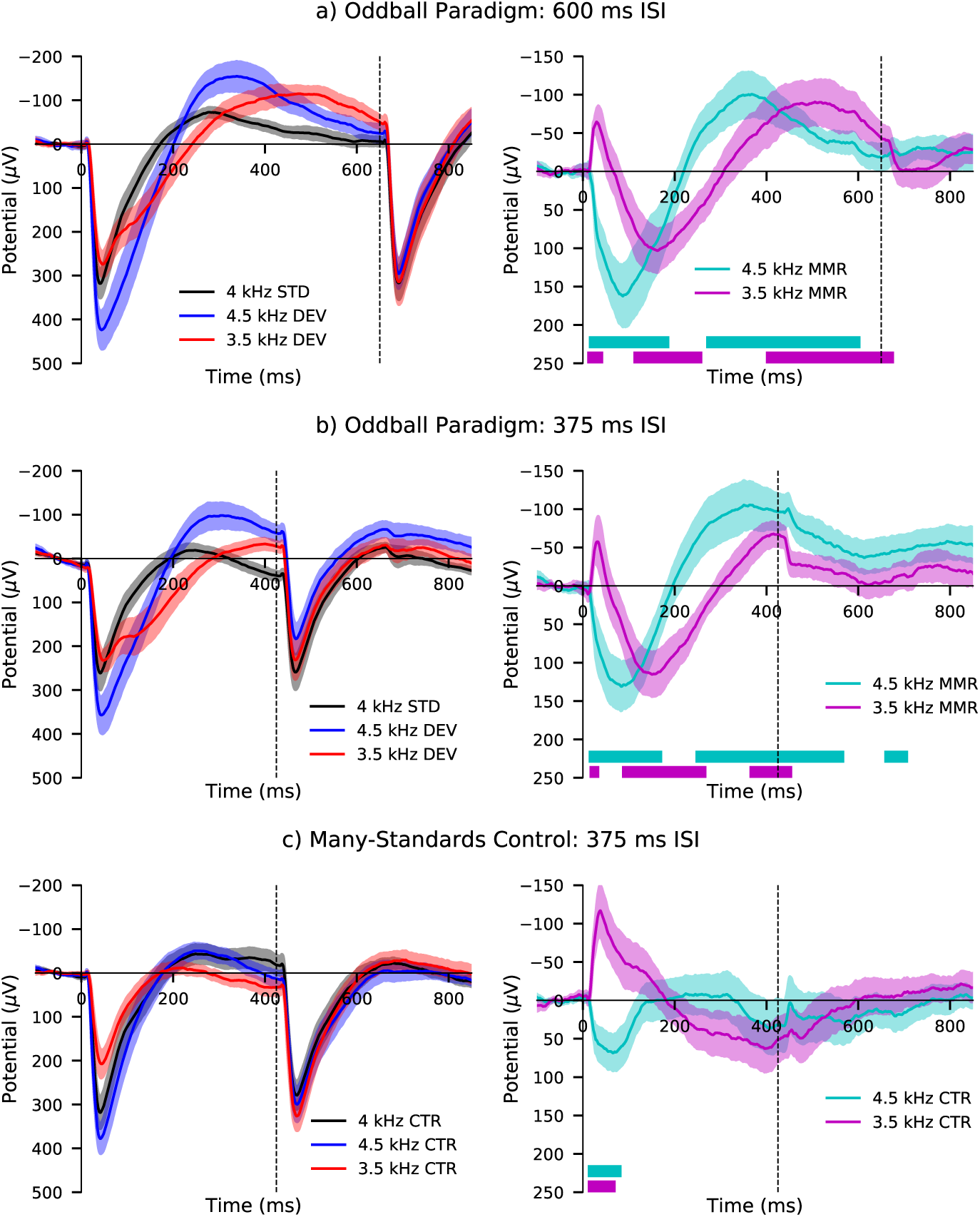
Auditory evoked potentials (left) and difference waveforms (right) from a) the oddball paradigm with 600 ms ISI; b) oddball paradigm with 375 ms ISI, and; c) many-standards control with 375 ms ISI. Note y-axes scale inversion. Onset times of second stimuli are annotated with vertical dashed lines. Regions of statistically significant (p < 0.05) deviation from zero in difference waveforms are annotated with solid blocks of the same colour as their respective traces. Comparing responses from (a) and (b) with those of (c), long-latency potentials over roughly 200 to 700 ms appear to be elicited by the oddball condition. It may also be noted that marginal differences exist between the responses to second stimuli in each set of AEP pairs.

### 2.3. Statistical Analysis

Wilcoxon signed-rank tests were performed on difference waveform data at each time point from 0 to 700 ms after stimulus onset, with the alpha value conventionally set to 0.05. This criterion was not adjusted for multiple-comparisons; such methods are arguably too conservative to appropriately evaluate these time-varying electrophysiology signals with high sampling rates. This statistical analysis was designed to identify latency ranges where difference waveforms significantly deviated from zero; the results are annotated in the respective panels on the right side of Figure 1.

### 2.4. Data Availability

The data used in this investigation is available from the originating researchers upon reasonable request (Kurkela et al., 2018).

## 3. Results

Auditory evoked potentials and their corresponding difference waveforms from both oddball paradigms and the many standards control paradigm are plotted in Figure 1. It can be seen by comparing difference waveform morphologies that both oddball paradigms appear to evoke pronounced activity extending through 200 to 700 ms. These long-latency potentials are conspicuously absent from control paradigm waveforms.

Early differences in many-standards control waveforms occurring before 100 ms may be related to differences in stimulus frequency, given the opposing polarities exhibited by ascending and descending frequency changes. Influences of different physical properties of stimuli on AEP and mismatch response waveforms measured from anaesthetised and conscious mice have been quite clearly demonstrated (O’Reilly and Conway, 2019). In contrast, mismatch responses from the oddball paradigm initially display opposite polarity trajectories, presumably due to their opposing frequency deviances, before becoming more coherent and displaying similar waveform morphologies, which may be attributed to their oddball context.

In light of this, it appears that the complete mismatch response might consist of two distinct components appearing after differences in stimulus onset responses caused by unequal frequency sensitivity. These two aspects of the deviant-induced response manifest here as a negative-going peak from 100 to 200 ms followed by a positive-going peak between 200 and 700 ms.

## 4. Discussion

Shorter latency components (<200 ms) observed from this dataset have been analysed already (Kurkela et al., 2018). The early effect of stimulus frequency on auditory onset responses was identified in the 30 to 70 ms window, whereas differences between standard and oddball conditions were found across the latency range of 80 to 180 ms. The authors deemed this to reflect a true rodent mismatch response, and the present analysis does not contradict this interpretation. Following the analysis in Figure 1, the 100 to 200 ms region may be characterised as the first component of the mismatch response, which displays positive peak amplitude here. This is immediately followed by a large negative amplitude deflection, peaking at approximately 300 to 500 ms post-stimulus. In the 375 ms ISI oddball paradigm, visualising this longer-latency component to a satisfactory conclusion is problematic without applying the double-epoch subtraction. Although a small amount of distortion appears to be caused by the onset response of second stimuli in each AEP pair, it is evident when comparing 375 ms and 600 ms ISI paradigm difference waveforms that both contain long-latency potentials with comparable trajectories. It may be noted that the double-epoch subtraction is not strictly required to observe long-latency MMR components from 600 ms ISI oddball paradigm waveforms if they are plotted up to the second stimulus onset at 650 ms.

Although not of identical polarity, these waveforms are considered to resemble those previously recorded from urethane-anaesthetised mice in response to frequency and increasing intensity deviant stimuli (O’Reilly, 2019). Waveform polarity differences may be caused by recording configuration, such as the position of ground and reference electrodes (Luck, 2014). As such, polarity cannot be relied upon for identifying alike components across studies that have applied differing conditions. The time-course of evoked activity may provide a more suitable means of identifying correspondences, providing there is also supporting rational. In this instance, both studies were performed in urethane-anaesthetised mice with frequency-varying oddball paradigms, demonstrating similar long-latency responses to deviant stimuli; therefore it may be said with some degree of confidence that these findings reflect comparable neurophysiological phenomena. There is also evidence of a late mismatch response in multi-unit recordings from the auditory cortex of anaesthetised mice, demonstrated by prolonged spiking activity approximately 200 to 400 ms after deviant stimulus presentation (Chen et al., 2015). The combined actions of large numbers of neurons contribute towards the epidural field potential, hence it is reasonable to expect many synchronised cell activations to occur during this latency range. Data from another recent study highlights increases in multi-unit activity recorded from prefrontal cortices in urethane-anaesthetised rats during frequency-varying auditory stimulation; displaying a long-latency response profile (200 to 700 ms) in response to oddball and deviant-alone paradigm stimuli (Supplementary Figure 1; Casado-Román et al., 2019). This supports the recently posited interpretation that oddball and deviant-alone conditions evoke similar neurophysiological responses (O’Reilly, 2019).

These findings collectively may pose a conundrum for the field of preclinical mismatch negativity research, considering that the time-course of this response extends outside of the accepted range of 100 to 250 ms for human MMN (Näätänen et al., 2012). This is exacerbated because rodent AEP components are thought to have shorter latencies than their human counterparts, due to having smaller brain sizes and therefore less distance for neurophysiological signals to travel (Siegel et al., 2003). Moreover, it has been shown that these long-latency mismatch responses are not elicited by quieter oddball stimuli (O’Reilly, 2019), which does not agree with some previous reports of human MMN being caused by quieter oddballs (Näätänen et al., 1989; Pakarinen et al., 2007). Careful research must be undertaken to determine the relationships between these waveforms. It is plausible that these long-latency potentials reflect mechanisms akin to a rodent P3a analogue. The P3a component in humans is reported to peak between 200 and 300 ms (Tavakoli et al., 2019). In contrast with the P3b, it does not require the subject’s conscious attention to be evoked, and is reportedly observed from patients who are asleep or anaesthetised (Koelsch et al., 2006; Plourde et al., 1993). Given these properties, the P3a may present a more likely equivalent to deviant-evoked long-latency potentials observed from anaesthetised rodents. This also agrees with the proposed *N1-MMN-P3a-RON* (reorienting negativity) pathway, whereby P3a is initiated downstream from MMN (Horváth et al., 2008); perhaps the earlier mismatch response component (100 to 200 ms) reflects human MMN, while the later mismatch response component (300 to 500 ms) reflects the human P3a. It is impossible to say definitively at this stage, and intricate work must be carried out to consider whether there are any tangible links between human and rodent AEPs.

## 5. Acknowledgements

We are very grateful for the openness and commitment to collective scientific endeavour demonstrated by Dr. Jari L. O. Kurkela and his colleagues at University of Jyväskylä, Finland. Without their willingness to share data openly this study would not have been possible.

## References

Baldeweg, T., Hirsch, S.R., 2015. Mismatch negativity indexes illness-specific impairments of cortical plasticity in schizophrenia: A comparison with bipolar disorder and Alzheimer’s disease. International Journal of Psychophysiology 95. doi:10.1016/j.ijpsycho.2014.03.008.

Casado-Román, L., Pérez-González, D., Malmierca, M.S., 2019. Prediction errors explain mismatch signals of neurons in the medial prefrontal cortex. [Pre-print] bioRxiv doi:10.1101/778928.

Chen, I.W., Helmchen, F., Lütcke, H., 2015. Specific Early and Late Oddball-Evoked Responses in Excitatory and Inhibitory Neurons of Mouse Auditory Cortex. The Journal of neuroscience: the official journal of the Society for Neuroscience 35, 12560–73. doi:10.1523/JNEUROSCI.2240-15.2015.

Harms, L., Michie, P.T., Näätänen, R., 2016. Criteria for determining whether mismatch responses exist in animal models: Focus on rodents. Biological Psychology doi:10.1016/j.biopsycho.2015.07.006.

Horváth, J., Winkler, I., Bendixen, A., 2008. Do N1/MMN, P3a, and RON form a strongly coupled chain reflecting the three stages of auditory distraction? Biological Psychology 79, 139–147. doi:10.1016/j.biopsycho.2008.04.001.

Koelsch, S., Heinke, W., Sammler, D., Olthoff, D., 2006. Auditory processing during deep propofol sedation and recovery from unconsciousness. Clinical Neurophysiology 117, 1746–1759. doi:10.1016/j.clinph.2006.05.009.

Kurkela, J.L., Lipponen, A., Kyläheiko, I., Astikainen, P., 2018. Electrophysio-logical evidence of memory-based detection of auditory regularity violations in anesthetized mice. Scientific Reports 8, 3027. doi:10.1038/s41598-018-21411-z.

Luck, S.J., 2014. An introduction to the event-related potential technique. MIT press.

Näätänen, R., Kujala, T., Escera, C., Baldeweg, T., Kreegipuu, K., Carlson, S., Ponton, C., 2012. The mismatch negativity (MMN)–a unique window to disturbed central auditory processing in ageing and different clinical conditions. Clinical Neurophysiology 123, 424–458.

Näätänen, R., Paavilainen, P., Alho, K., Reinikainen, K., Sams, M., 1989. Do event-related potentials reveal the mechanism of the auditory sensory memory in the human brain? Neuroscience Letters 98, 217–221. doi:10.1016/0304-3940(89)90513-2.

Nakamura, T., Michie, P.T., Fulham, W.R., Todd, J., Budd, T.W., Schall, U., Hunter, M., Hodgson, D.M., 2011. Epidural auditory event-related potentials in the rat to frequency and duration deviants: evidence of mismatch negativity? Frontiers in psychology 2, 367.

O’Reilly, J.A., 2019. Double-epoch subtraction reveals long-latency mismatch response in urethane-anaesthetized mice. Journal of Neuroscience Methods 326, 108375. doi:10.1016/j.jneumeth.2019.108375.

O’Reilly, J.A., Conway, B.A., 2019. Classical and controlled auditory mismatch responses to multiple physical deviances in anaesthetised and conscious mice. [Pre-print] bioRxiv doi:10.1101/831016.

Pakarinen, S., Takegata, R., Rinne, T., Huotilainen, M., Näätänen, R., 2007. Measurement of extensive auditory discrimination profiles using the mismatch negativity (MMN) of the auditory event-related potential (ERP). Clinical Neurophysiology 118, 177–185.

Plourde, G., Joffe, D., Villemure, C., Trahan, M., 1993. The P3a Wave of the Auditory Event-related Potential Reveals Registration of Pitch Change during Sufentanil Anesthesia for Cardiac Surgery. Anesthesiology: The Journal of the American Society of Anesthesiologists 78, 498–509.

Ruusuvirta, T., Penttonen, M., Korhonen, T., 1998. Auditory cortical event-related potentials to pitch deviances in rats. Neuroscience letters 248, 45–48.

Siegel, S.J., Connolly, P., Liang, Y., Lenox, R.H., Gur, R.E., Bilker, W.B., Kanes, S.J., Turetsky, B.I., 2003. Effects of strain, novelty, and NMDA blockade on auditory-evoked potentials in mice. Neuropsychopharmacology doi:10.1038/sj.npp.1300087.

Tavakoli, P., Dale, A., Boafo, A., Campbell, K., 2019. Evidence of P3a During Sleep, a Process Associated With Intrusions Into Consciousness in the Waking State. Frontiers in Neuroscience 12, 1028. doi:10.3389/fnins.2018.01028.

